# A separation-of-function *ZIP4* wheat mutant allows crossover between related chromosomes and is meiotically stable

**DOI:** 10.1101/2021.09.03.458546

**Authors:** Azahara C. Martín, Abdul Kader Alabdullah, Graham Moore

## Abstract

Many species, including most flowering plants, are polyploid, possessing multiple genomes. During polyploidisation, fertility is preserved via the evolution of mechanisms to control the behaviour of these multiple genomes during meiosis. On the polyploidisation of wheat, the major meiotic gene *ZIP4* duplicated and diverged, with the resulting new gene *TaZIP4-B2* being inserted into chromosome 5B. Previous studies showed that this *TaZIP4-B2* promotes pairing and synapsis between wheat homologous chromosomes, whilst suppressing crossover between related (homoeologous) chromosomes. Moreover, in wheat, the presence of *TaZIP4-B2* preserves up to 50% of grain number. The present study exploits a ‘separation-of-function’ wheat *Tazip4-B2* mutant named *zip4-ph1d*, in which the *Tazip4-B2* copy still promotes correct pairing and synapsis between homologues (resulting in the same pollen profile and fertility normally found in wild type wheat), but which also allows crossover between the related chromosomes in wheat haploids of this mutant. This suggests an improved utility for the new *zip4-ph1d* mutant line during wheat breeding exploitation, compared to the previously described CRISPR *Tazip4-B2* and *ph1* mutant lines. The results also reveal that loss of suppression of homoeologous crossover between wheat chromosomes does not in itself reduce wheat fertility when promotion of homologous pairing and synapsis by *TaZIP4-B2* is preserved.

## Introduction

Polyploidy exists in a wide range of species including yeast, flowering plants, fish, flatworms, shrimp and amphibians [1,2,3,4], and in two different forms, namely allopolyploidy and autopolyploidy. Allopolyploids, such as tetraploid (pasta) and hexaploid (bread) wheat, arise by combining related but not completely identical genomes, whereas autopolyploids arise via multiplication of a basic set of chromosomes. Previous studies have revealed extensive chromosome rearrangements in some polyploids leading to changes in gene content and/or expression [3,5,6,7]. However, recent large-scale genome sequencing and RNA analysis has revealed that homoeologous (related) chromosomes of hexaploid wheat (*Triticum aestivum* L.) did not exhibit extensive gene loss or expression changes following polyploidisation [8,9]. This suggests that a major factor rapidly evolved upon wheat polyploidisation to control the behaviour of its multiple genomes during meiosis, hence preserving fertility. It is generally accepted that in wheat, this control is due to an event that occurred on chromosome 5B following polyploidisation [10,11].

Hexaploid wheat lines lacking the whole of chromosome 5B, were used to generate haploids and wheat-wild relative hybrids, both of which exhibited chiasmata (physical attachments at which crossover and recombination occur) between homoeologous chromosomes at metaphase I [10,11]. Indeed, the occurrence of this crossover (CO) between the homoeologous chromosomes of wheat and its wild relatives in such hybrids was a key observation, allowing their subsequent use for the introgression of desirable chromosome segments from wild relatives into wheat during breeding. Sears [12] identified a 5B deletion mutant (now known to be 59.3Mb in size [13]) termed *ph1b*, which has been used for this purpose over the last 60 years. However, this line also showed meiotic abnormalities and reduced fertility, leading Riley and others [10] to propose that the suppression of homoeologous CO between wheat chromosomes observed in wheat haploids (where only homoeologous chromosomes are present), was important for stabilising the wheat genome and preserving its fertility. It had therefore become generally accepted that homoeologous CO suppression was necessary for polyploid stability.

It was unclear from these studies whether the controlling factor on chromosome 5B was due to a single or multiple genes. At that time, the term ‘pairing’ was used to describe the occurrence of chiasmata between chromosomes at metaphase I. Thus, the term ‘pairing homoeologous’ (*Ph1*) subsequently became the accepted term for the chromosome 5B associated ‘suppression of homoeologous CO’ phenotype observed in haploids and wheat-wild relative hybrids [14]. Currently however, the term ‘pairing’ is used to describe the initial association of meiotic chromosomes prior to synapsis, which is how it will be subsequently used in this paper.

In wild type wheat, telomeres cluster as a bouquet at one nuclear pole during early meiosis, and homologous chromosomes (homologues) pair and synapse from these telomere regions. In wild type wheat-rye hybrids (where no homologues are present), homoeologous chromosomes (homoeologues) can also synapse; however, synapsis only occurs after the telomere bouquet stage, when telomeres have dispersed [15]. Synaptonemal complex studies also revealed that a ‘promotion of homologous synapsis’ phenotype was also associated with *Ph1* on chromosome 5B [15,16,17]). Homologous chromosome sites were pairing early at the telomere bouquet stage in wild type wheat, which led to the promotion of homologous synapsis, reducing the chances of homoeologous synapsis occurring after the telomere bouquet stage [15]. Therefore, although *Ph1* was originally defined as a ‘suppressing homoeologous CO’ phenotype, a ‘promotion of pairing-synapsis’ phenotype was also associated with the 59.3Mb region defined by the *ph1b* deletion. As a consequence, the wheat *ph1b* mutant exhibited major meiotic abnormalities (univalents and/or multivalents) in around 56% of meiocytes [18].

Both the ‘homoeologous CO suppression’ and the ‘promotion of homologous pairing-synapsis’ phenotypes were defined to the same region of chromosome 5B, possessing a copy of the major meiotic gene *ZIP4* (*TaZIP4-B2*) that duplicated and diverged from chromosome 3B [19,20]. Thus, hexaploid wheat carries four copies of *ZIP4*, one copy on each of the group 3 chromosomes and a fourth copy, the duplicated and diverged *TaZIP4-B2* on chromosome 5B. *ZIP4* has previously been shown to be required for 85% of homologous COs in yeast, Sordaria, mammals, rice and Arabidopsis, and for the initial pairing and synapsis in yeast and Sordaria [21,22,23,24]. Thus, although in *Arabidopsis* and rice, *ZIP4* has only been shown to be required for homologous CO and not synapsis, in yeast, *ZIP4* is required for both CO and synapsis. Analysis of a CRISPR deletion mutant of *TaZIP4-B2* and of the Sears *ph1b* deletion mutant, revealed the presence of major meiotic abnormalities in around 56% of meiocytes. Both mutants also revealed a similar level of CO between homoeologous chromosomes when crossed with the same wild relative [25]. This confirms that the duplicated and diverged *TaZIP4-B2* copy is responsible for both the ‘suppression of homoeologous CO’ and the ‘promotion of homologous pairing-synapsis’ phenotypes defined on chromosome 5B. Moreover, the presence of this duplicated copy (*TaZIP4-B2*) has also recently been shown to be necessary for preservation of 50% of grain number [26].

Early cytogenetic studies suggested that the ‘suppression of homoeologous CO’ phenotype is required for genome stability and preservation of wheat fertility. However, it is also possible that the effect of *TaZIP4-B2* on promotion of homologous pairing and synapsis is relevant or even more important. The availability and analysis of a separation-of-function *Tazip4-B2* mutant named *zip4-ph1d* (formerly TILLING line *Tazip4-B2*-Cadenza1691 [20]) has allowed us to investigate this further, confirming that indeed, suppression of homoeologous CO by *ZIP4* is not essential for wheat fertility.

## Results and Discussion

The CRISPR *Tazip4-B2* mutant showing complete loss of the TaZIP4-B2 function phenotype carries a deletion of 38 amino acids (A^104^ to E^141^) that encompasses the 1^st^ TPR (tetratricopeptide repeat) in the highly conserved SPO22 domain [25,26]. We previously identified another mutant, a *Tazip4-B2*-Cadenza1691 TILLING mutant, which carries a single missense mutation (C to T change at position 500 in the coding sequence (CDs), leading to an amino acid substitution A^167^V) within the same SPO22 domain [20]. When wild type wheat was crossed with the wild relative *Aegilops variabilis*, very few homoeologous COs were observed in the resulting hybrid. However, when either of the two mentioned *Tazip4-B2* mutants were crossed with the same wild relative, an increased level of homoeologous CO was observed, similar to that found when the original Sears 59.3Mb *ph1b* deletion mutant was crossed with this same wild relative [20,25]. This indicates that in both mutants, the capability of *TaZIP4-B2* to suppress CO between homoeologous wheat and wild relative chromosomes has been lost. However, there was a small but very important phenotypic difference between the two mutants. In the CRISPR *Tazip4-B2* mutant, 56% of meiocytes had major abnormalities, with multivalents present in 32% meiocytes (average 0.39 per meiocyte). However, in the *Tazip4-B2*-Cadenza1691 mutant, only 26% of meiocytes analysed had abnormalities, with no multivalents present [20]. These results suggest that while the Cadenza1691 *TaZIP4-B2* mutant has lost the function to suppress homoeologous CO, it has retained the ability of *TaZIP4-B2* to promote pairing-synapsis between homologues. Thus, the Cadenza1691 *TaZIP4-B2* mutant represents a ‘separation-of-function’ *TaZIP4-B2* mutant. We have since renamed this line as the *zip4-ph1d* mutant and further analysed its phenotype.

### The pollen profiles of the *zip4-ph1d* mutant and wild type wheat are the same

A previously developed pollen profiling approach revealed that 50% of CRISPR *Tazip4-B2* mutant pollen grains were small and unviable, correlating with 56% meiocytes possessing major meiotic abnormalities [26]. In contrast, major disruption of pollen development was considered less likely in the new *zip4-ph1d* mutant, as it lacks major meiotic abnormalities, with only some meiocytes showing occasional univalents. The pollen profiling method was thus performed to compare pollen grain size distribution and number from the *zip4-ph1d* mutant with that from wild type wheat. Pollen was collected from full mature anthers (just before opening) from mutant and wild type wheat (*T. aestivum* cv. Cadenza). Five samples were collected from main florets in the middle portion of the first spike of each plant. Each sample contained three anthers belonging to the same floret. Pollen grain size and number were measured from 50 samples of each genotype using the Coulter counter Multisizer 4e. In this study, a mean of 8490 pollen grains were measured from each genotype (average 902±34 and 796±29 pollen grains per mutant and wild type plant respectively) (Table 1).

**Table 1.** Summary of pollen analysis data for *zip4-ph1d* mutant and wild type (WT) wheat Cadenza.

| Genotype | Total measured pollen grains | Measured pollen grains per plant ± SE | Pollen size (µm) ± SE | Pollen number per anther ± SE | Small pollen grain (%) ± SE | Pollen viability % ± SE |
| --- | --- | --- | --- | --- | --- | --- |
| *zip4-ph1d* | 9017 | 901.69±4.44 | 47.64±0.50 | 3005.64±114.80 | 15.00±2.14 | 95.39±1.20 |
| Cadenza WT | 7963 | 796.26±29.19 | 47.57±0.51 | 2654.2±97.31 | 14.04±1.56 | 96.02±0.51 |

Surprisingly, the *zip4-ph1d* mutant displayed the same single major peak pollen profile as the wild type (Fig. 1A), with average pollen sizes of 47.64±10.50 µm and 47.57±0.51 µm, in the mutant and wild types respectively. Conversely, the CRISPR *Tazip4-B2* mutant displayed a two peak pollen profile [26], with the first peak corresponding to pollen grains with grain size distribution similar to that of both the wild type and *zip4-ph1d* pollen, and the second peak corresponding to pollen with smaller grain size. Thus, the *zip4-ph1d* mutant did not produce significant numbers of smaller size pollen grains (15±2.14%, comparable to that of wild type 14.04±1.56%) (Fig. 1C), in contrast to the percentage of small pollen produced by the CRISPR *Tazip4-B2* (and *ph1b)* mutant plant (*ca*. 50%) [26]. The average number of pollen grains per anther of the *zip4-ph1d* mutant (3006±115) was slightly higher than that of the Cadenza wild type (2654±97), however this difference was not significant at *p*-value < 0.01. (Fig. 1B).

**Figure 1.**
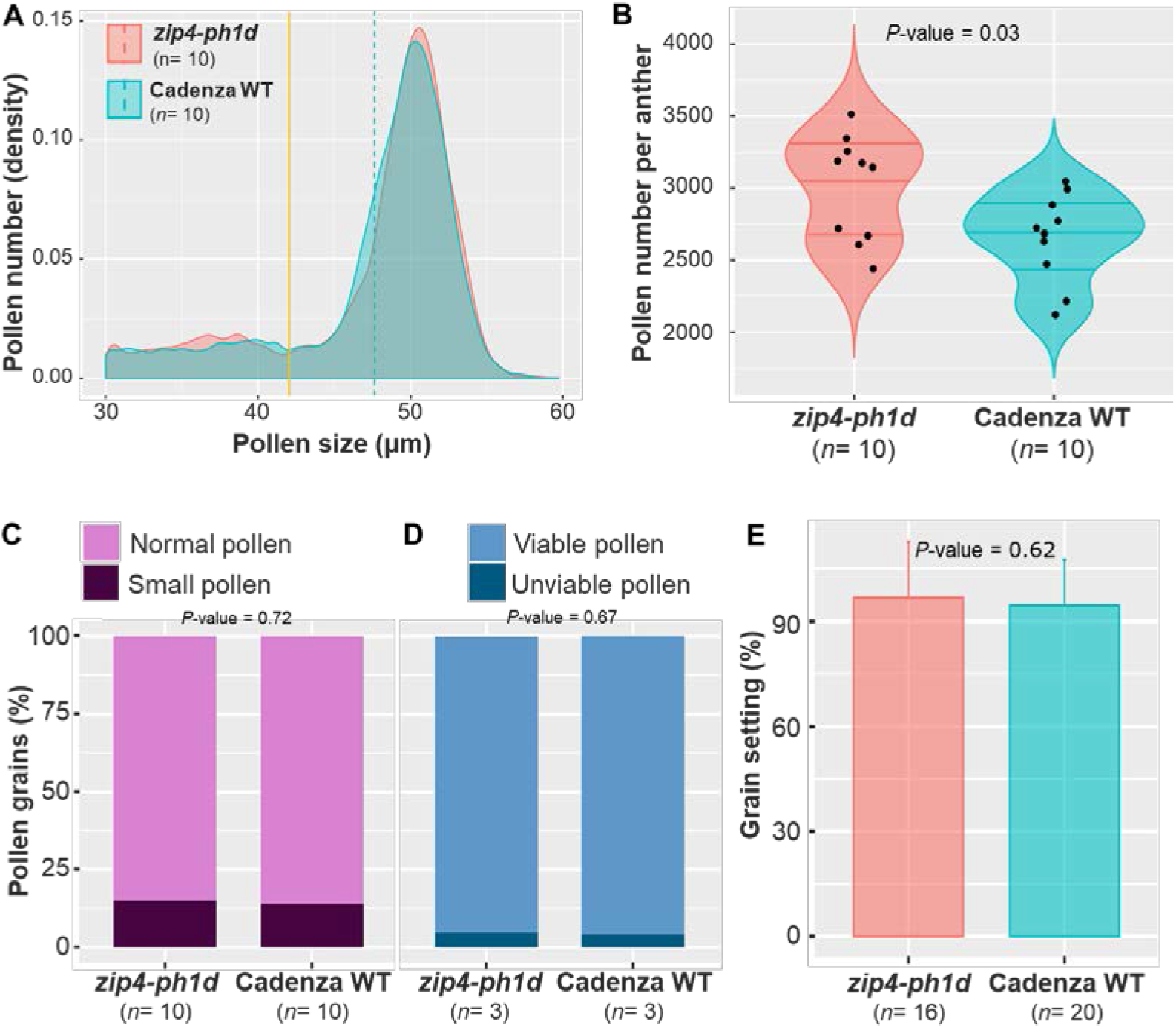
Pollen and grain setting analysis of the *zip4-ph1d* mutant and its wild type. (**A**) Density plot of differential pollen size distribution data collected by coulter counter (Multiziser 4e) showing similar pollen profiles in the *zip4-ph1d* mutant and its wild type (cv. Cadenza). Dotted lines indicate mean pollen grain size for each genotype. Yellow lines indicate the borderline between normal and small pollen for each genotype. (**B**) Violin plot showing number of pollen grains per anther in the *zip4-ph1d* mutant and its wild type. No significant differences in pollen number per anther were found between any of the mutants and their wild types. (**C**) Percentages of small pollen grains in each genotype. Pollen grains ≤42 µm are considered small. (**D**) Percentage pollen viability for each genotype, according to the Alexander staining method. More than 3000 pollen grains per genotype were scored after Alexander staining. (**E**) Grain setting (normalised grain number per spike) in the *zip4-ph1d* mutant and wild type control (cv. Cadenza) under CER growth conditions. *n* refers to the number of biological replicates used in each experiment. Error bar refers to the standard deviation.

We also assessed the viability of pollen from the *zip4-ph1d* mutant using Alexander staining. Previous studies had revealed that the *CRISPR-Tazip4-B2* and *ph1b* mutants resulted in similar percentages of unviable pollen grains, being 28% and 25.8% respectively [26], while unviable pollen in the corresponding wild type wheat did not exceed 3.3% on average. More than 3000 pollen grains from the *zip4-ph1d* mutant and Cadenza WT were scored (from three biological replicates per genotype), after Alexander staining and image acquisition. Pollen coloured dark magenta after treatment with Alexander stain was considered viable, whereas light blue-green stained pollen was considered unviable. Consistent with our pollen profiling analysis, unviable pollen grains in the *zip4-ph1d* mutant did not exceed 5% like in wild type wheat (Fig. 1D), suggesting again that the pollen in the *zip4-ph1d* mutant is perfectly normal.

### Grain setting is not reduced in the *zip4-ph1d* mutant

The absence of any major meiotic abnormality in most *zip4-ph1d* mutant meiocytes, in conjunction with a pollen profile similar to that found in wild type wheat, suggested that the *zip4-ph1d* mutant should be mainly fertile. To verify this, grain setting analysis was performed on both the *zip4-ph1d* mutant and wild type wheat. Spikelet number was recorded, as well as the number of grains per spike for the first three spikes from the mutant and wild type plants. The normalized grain number per spike was used to compare genotypes. The experiment, involving over 15 plants (biological replicates) for each genotype, confirmed that there was no significant difference (*p*-value = 0.62) observed in seed set between the *zip4-ph1d* mutant (96.76±3.98%) and wild type wheat (94.36±2.95%) (Fig. 1E). This result was markedly different from the fertility of the CRISPR *Tazip4-B2* mutant which exhibited up to a 50% reduction in grain number. Thus, the increased fertility of the *zip4-ph1d* mutant, compared to the CRISPR *Tazip4-B2* mutant, is consistent with *zip4-ph1d* being a partial loss-of-function mutant.

### Effect of the A^167^V substitution resulting in partial loss-of-function

ZIP4 is a tetratricopeptide repeat (TPR) containing protein that has been shown to interact with chromosome axis and crossover formation proteins in budding yeast (*Saccharomyces cerevisiae*) [27]. Studies of other proteins containing TPRs have shown that they form alpha solenoid helix structures which play an important role in the formation of multiple protein-protein binding interactions and facilitate the assembly of proteins into complexes [28,29]. Our previous study showed that half of the wheat ZIP4 protein is composed of TPRs [27]. Within this context, we performed an *In Silico* protein sequence analysis to gain insight into why the amino acid substitution (A^167^V) within TaZIP4-B2 resulted in its partial loss-of-function in the *zip4-ph1d* mutant. Multiple amino acid sequence alignments were conducted to assess the level of amino acid sequence conservation between ZIP4 proteins from different organisms. A total of 45 protein sequences from 38 species belonging to a wide range of organisms including plants (monocots and dicots), animals (mammals, birds, and fishes) and budding yeast were included in the analysis. Strikingly, the alanine residue (A167) substituted in *zip4-ph1d* was found highly conserved in all aligned sequences (Fig. 2A), indicating the potential importance of this amino acid in either the function and/or structure of the protein. We previously reported that TaZIP4 wheat proteins possess 12 TPR domains, with the TaZIP4-A1 protein diverging in the TPR10, TPR11 and TPR12 regions, and the TaZIP4-B2 protein diverging in the TPR3 region, such that these domains are no longer identified as TPRs [26]. The TPR consensus sequence contains 34 residues, in which the conserved positions are W4-L7-G8-Y11-**A20**-F24-A27-P32 [30]. The alanine residue substituted in the *zip4-ph1d* mutant is precisely the one located at the conserved position 20 of the TPR2 in the TaZIP4-B2 copy.

**Figure 2.**
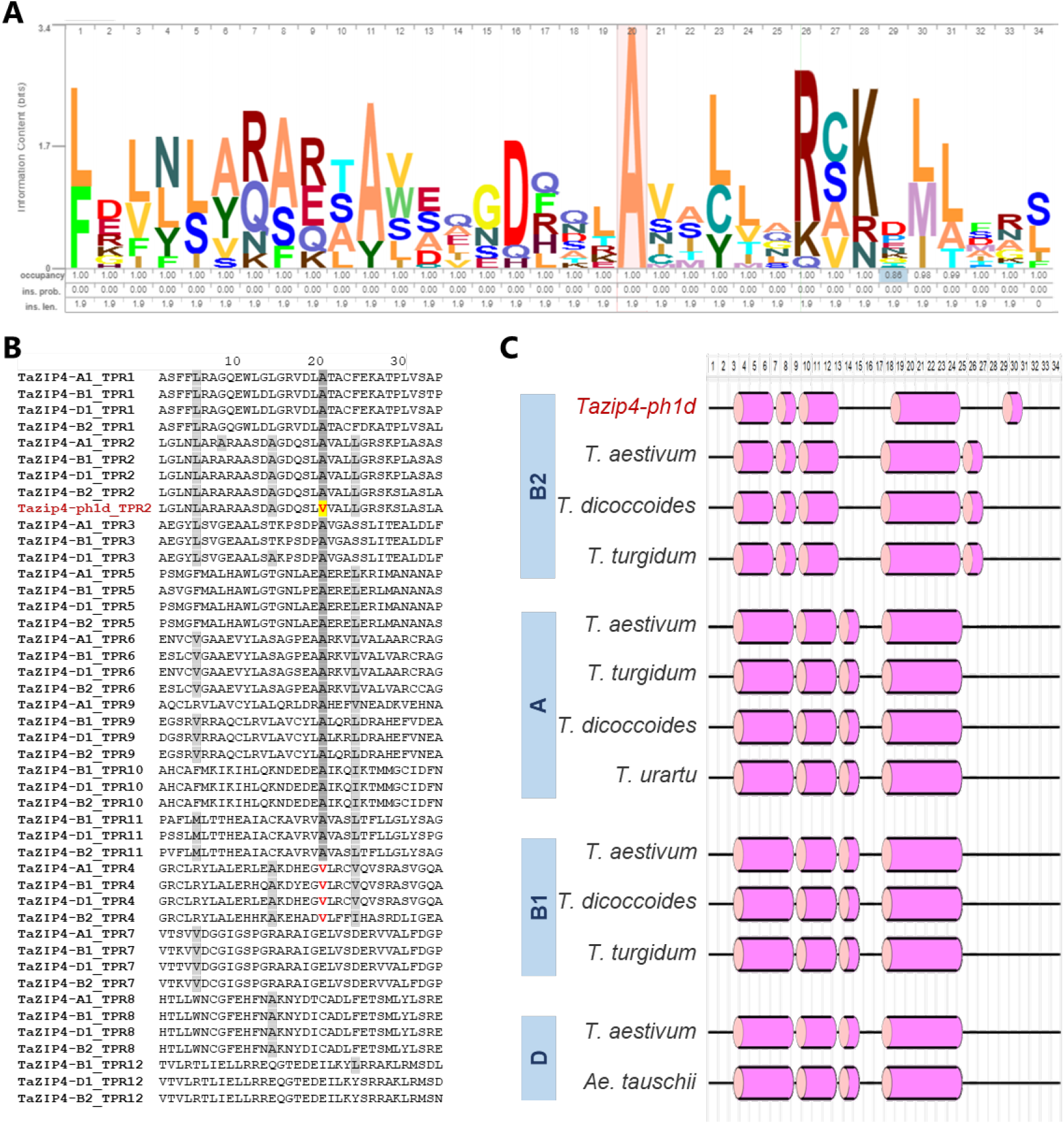
ZIP4 protein sequence analyses. (**A**) Graphic representation of the conserved residues of the TaZIP4-B2 TPR2 domain and the corresponding sequences from a wide range of organisms including plants, animals and yeast. (**B**) Peptide sequence comparison of the TPR domains in the four TaZIP4 proteins. (**C**) Predicted secondary structure of the TPR2 domain in the ZIP4 protein of the *Tazip4-ph1d* mutant, compared with those of hexaploid and tetraploid wheat, and their progenitors. The secondary structure was predicted using the PSSM random forest-based model in the PEP2D server. Helix is represented by the pink cylindrical shape and coil is represented as a black line.

Comparison of the peptide sequences of all TPRs in the four wheat TaZIP4 proteins (TaZIP4-A1, TaZIP4-B1, TaZIP4-B2 and TaZIP4-D1), revealed that there was an alanine residue in position 20 in the TPR1, TPR2, TPR3, TPR5, TPR6 and TPR9 regions of all the TaZIP4 proteins (Fig. 2B). Interestingly, a valine (V) residue was also found at position 20 in the TPR4 of all four wheat TaZIP4 proteins, which was the same amino acid substituted for alanine at position 20 within the TPR2 of TaZIP4-B2 within the *Tazip4-ph1d* mutant (Fig. 3B). This observation, together with the fact that both alanine and valine residues have very similar properties (both being small, aliphatic (non-polar) and hydrophobic amino acids), may explain why the A^167^V substitution in TaZIP4-B2 results in the protein still being partially functional, despite a substitution of the most conserved ZIP4 amino acid residue.

**Figure 3.**
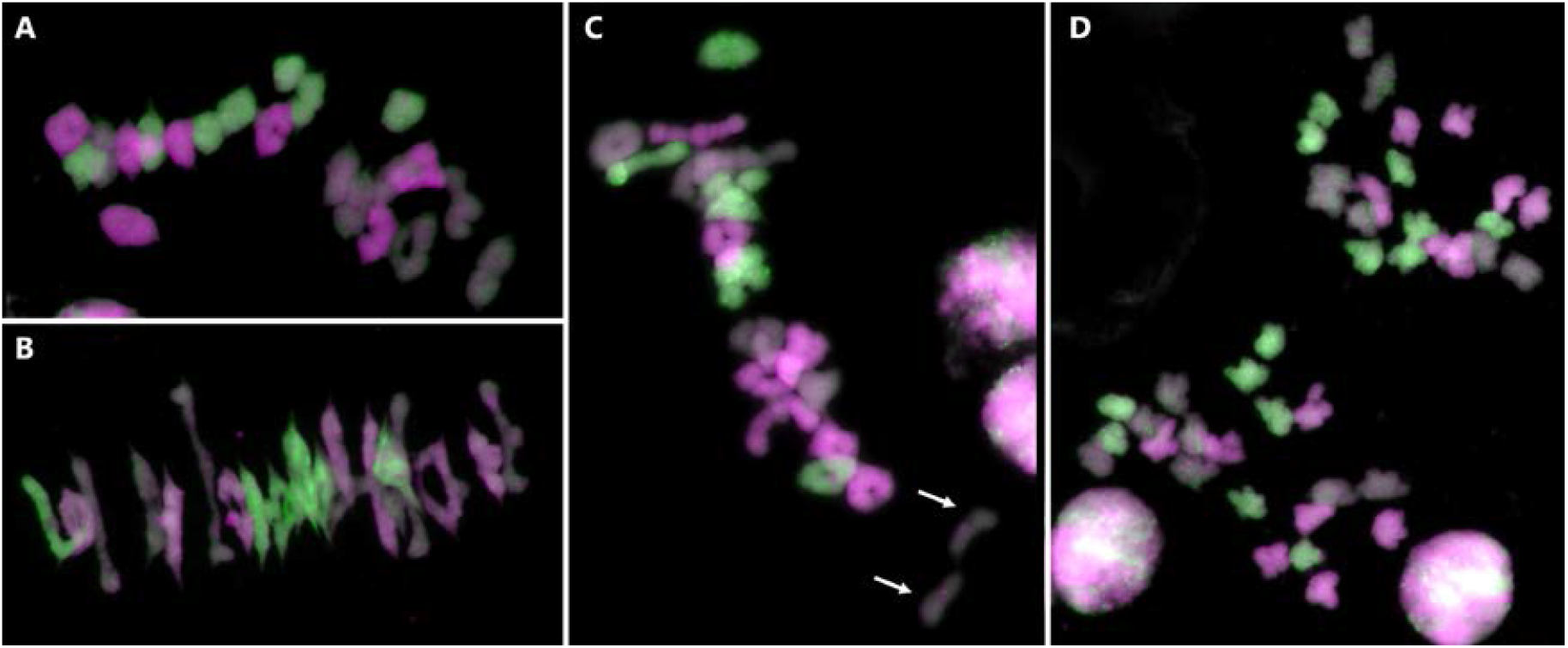
Genomic in *situ* hybridization (GISH) of meiotic metaphase I (A, B, C) and anaphase I (D) spreads in the *zip4-ph1d* mutant. A-genome chromosomes are shown in magenta, B-genome chromosomes in grey and D-genome chromosomes in green. (**A**) Representative metaphase I configurations showing 21 ring bivalents, (**B**) metaphase I showing 16 ring and 5 rod bivalents, and (**C**) metaphase I with 16 ring bivalents, 4 rod bivalents and 2 univalents indicated by an arrow. In (**D**), a representative anaphase I showing correct chromosome segregation.

Since the A>V substitution might have resulted in a subtle conformational change in the helical structure of TPR2, the 2D structures of TPR2s in the TaZIP4s derived from hexaploid wheat (including *Tazip4-ph1d*), tetraploid wheat and their progenitors were predicted and compared. The helical structure of the TPR2 in wild type TaZIP4-B2 (encoded by the chromosome 5B gene), was different from the TPR2 structures in ZIP4s encoded by the chromosome group 3 genes (*TaZIP4-A1, TaZIP4-B1* and *TaZIP4-D1*) (Fig. 3C). This may help to explain why the duplicated and diverged TaZIP4-B2 copy has a different function to the group 3 ZIP4s proteins [26]. Moreover, the presence of a valine at position 20 within its TPR2 results in a subtle difference in its helical structure, when compared to the structure of the TPR2 wild type possessing an alanine at the same location (Fig. 2C). Thus, the separation-of-function TaZIP4-B2 phenotype may result from a subtle conformational change in the TPR2 domain.

### Chiasmata take place between homologous chromosomes in the *zip4*-*ph1d* mutant

Previous studies using the CRISPR *Tazip4-B2* deletion mutant showed that some chiasmata occurred between wheat homoeologues at metaphase I, indicating that the TaZIP4-B2 protein (carrying an in-frame 38 amino acid CRISPR deletion (A^104^ to E^141^)) had lost the ability to both promote homologous pairing-synapsis and to suppress homoeologous CO [25]. However, the *zip4-ph1d* mutant (originally *Tazip4-B2*-Cadenza1691) showed an average of 17.42 ring bivalents, 3.29 rod bivalents and 0.58 univalents at meiotic metaphase I, with no multivalents present [20]. Moreover, 74% of the *zip4-ph1d* meiocytes showed no univalents, while 26% exhibited 2 univalents. We performed GISH on meiotic metaphase I spreads derived from the *zip4*-*ph1d* mutant, in order to assess whether the observed ring and rod bivalents were composed of homologues, and whether the univalents resulted from chiasmata failure between homologues. We analysed 328 meiocytes at metaphase I, derived from three biological replicates of *zip4-ph1d* mutants (more than 100 meiocytes from each replicate) and found only 4 meiocytes with homoeologous chiasmata (a single multivalent was present in each case) (Fig. 3A, B). This result contrasts with observations from the CRISPR *Tazip4-B2* mutant, where over 32% meiocytes had at least one multivalent (and hence homoeologous chiasmata). Previous studies have shown that even in haploids derived from wild type wheat, there are 0.7-1.5 chiasmata between wheat homoeologues per meiocyte [31]. This low level of homoeologous chiasmata at metaphase I in wild type wheat could be contributing to the abnormal pollen observed in wild type wheat [26]. More importantly however, in the *zip4*-*ph1d* mutant, the great majority of chiasmata occur between homologues, confirming that, as in wild type wheat, most chromosomes pair and CO correctly.

In the *zip4-ph1d* mutant, univalents at metaphase I are present in 26% of meiocytes, with most univalent pairs also being homologues (Fig. 3C). This confirms that the *Tazip4-B2* copy containing the A^167^V substitution retains the ability to promote pairing-synapsis between homologues. However, the relatively low level of univalents observed suggests that the *TaZIP4-B2* copy only have a small effect on homologous CO in wheat. Thus, it seems more likely that the *TaZIP4* copies on group 3 chromosomes are predominantly responsible for the homologous CO promotion, as observed in other species [21,22,23,24]. Surprisingly, the univalent pairs observed at metaphase I in the *zip4-ph1d* mutant were frequently close to each other and orientated to different poles of the nucleus (Fig. 3C). This suggests that they could have been connected to each other, as they were orientated correctly on the metaphase I plate, but there was CO failure at this later stage. Since the TaZIP4-B2 copy in the *zip4-ph1d* mutant can still promote homologous pairing-synapsis, it seems likely that these chromosomes had correctly synapsed and the recombinational machinery had been loaded, but CO formation had subsequently failed at the very last stage. This observation is consistent with a previous study [32] where, even if the meiotic protein MLH1 (which marks sites that will become COs) was loaded onto the paired wheat chromosomes at diplotene, some of the sites still failed to progress to COs at this late stage.

### There is mostly balanced chromosome segregation in the *zip4*-*ph1d* mutant

Homologous chromosomes connected by at least one chiasma at metaphase I, can be correctly segregated to a different nuclear pole at anaphase I. A second cell division leads to the subsequent formation of tetrads with four balanced gametes. In the CRISPR *Tazip4-B2* mutant, the presence of univalents and multivalents led to lagging chromosomes and split sister chromatids at anaphase I, with a consequently high percentage of tetrads presenting micronuclei (more than 50%) and a negative effect on pollen formation and fertility [26]. However, most metaphase I meiocytes in the *zip4*-*ph1d* mutant do not exhibit such major meiotic abnormalities, and pollen profile and fertility are the same as in wild type wheat.

Analysis of micronuclei level at the tetrad stage is a reliable approach to assess chromosome segregation problems during meiosis [26,33]. Therefore, tetrad analysis was undertaken to confirm that the *zip4*-*ph1d* mutant does not exhibit major chromosome segregation problems. We analysed tetrads from three biological replicates of each of the *zip4-ph1d* mutant and wild type Cadenza, with more than 700 tetrads scored from each genotype (Fig. 4H, Supplementary Table S1). As expected, no major disruption of chromosome segregation was detected in the mutant, as compared to that observed in the wild type Cadenza. The *zip4-ph1d* mutant was associated with 12.2% of tetrads with micronuclei, a small increase compared to that observed in wild type Cadenza (5.6%) (Fig. 4E-G), but a considerably lower number than the 50.6% of tetrads with micronuclei previously observed in the *CRISPR Tazip4-B2* deletion mutant [26] (Fig. 4G).

**Figure 4.**
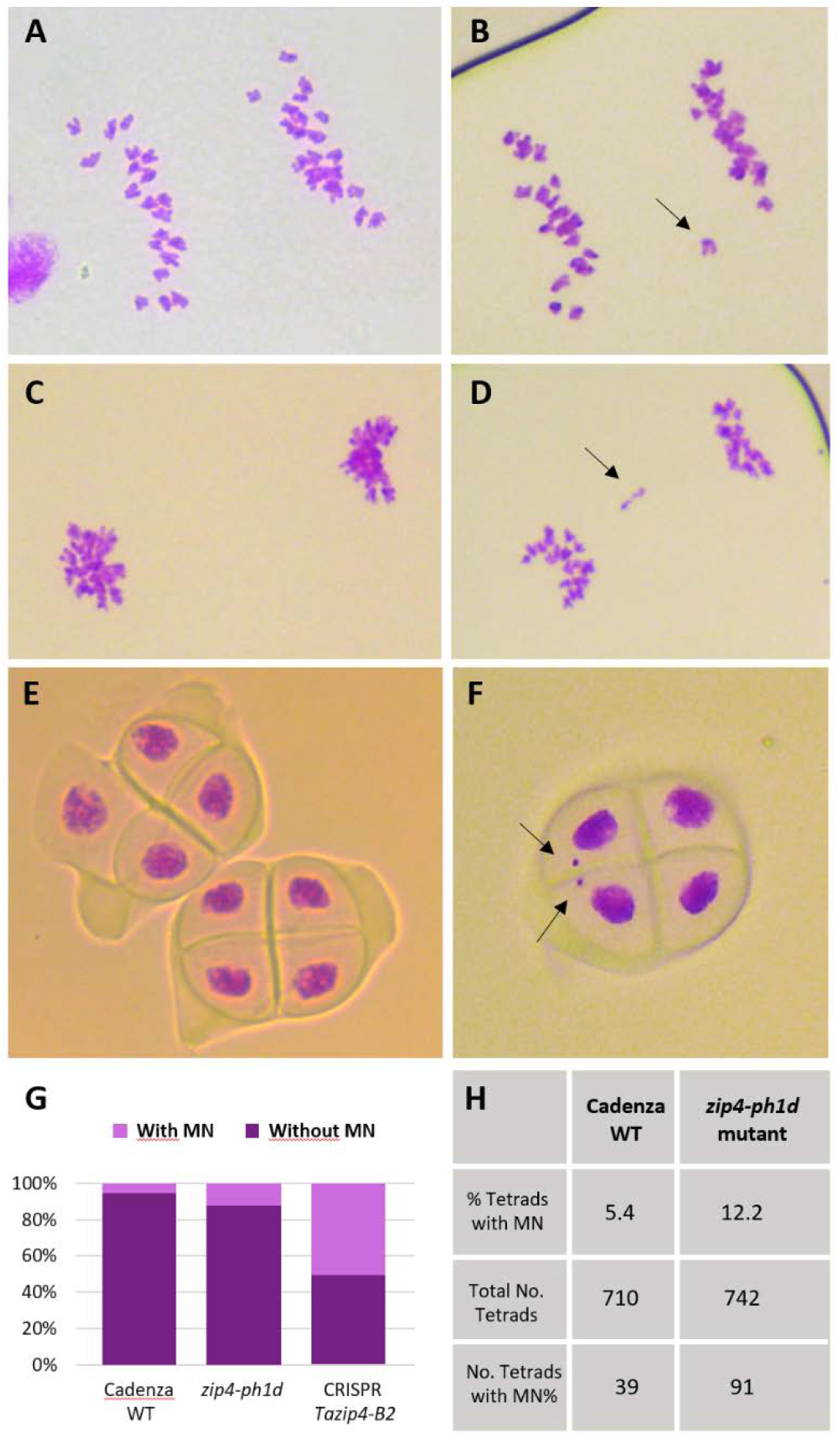
Chromosome segregation during meiosis in the *zip4-ph1d* mutant is mostly balanced. (**A, C**) Early and late anaphase I, displaying equal separation of homologous chromosomes to each pole of the nucleus, as observed in most meiocytes in the *zip4-ph1d* mutant and (**B, C**) Same meiotic stages showing lagging chromosomes at the metaphase plate indicated by an arrow. (**E**) Two perfectly formed tetrads in the *zip4-ph1b* mutant and (**F**) one tetrad with two micronuclei indicated by an arrow. (**G**) Comparison of tetrad percentage with micronuclei (MN) in wild type (WT) Cadenza, the *zip4-ph1d* mutant and the CRISPR *zip4-B2* mutant [26]. (**H**) Percentages of tetrads with and without MN in WT Cadenza and the *zip4-ph1d* mutant.

In the *zip4*-*ph1d* mutant, univalents were present in 26% meiocytes, resulting in 12.2% tetrads with micronuclei. Conversely, in the CRISPR *Tazip4-B2* mutant, 56% meiocytes with major abnormalities (32% with multivalents and 24% with univalents), leading to 50.6% tetrads with micronuclei, were reported previously [26]. These different scores suggest that more pairs of univalents are correctly segregated in the *zip4*-*ph1d* mutant, even though formation of chiasmata have not occurred. We obtained further confirmation of the correct segregation of these univalents, by analysing the meiotic configurations at anaphase I for the presence of laggard chromosomes. The majority of anaphases analysed lacked lagging chromosomes (Fig. 4A,C), with only 8.6% of them showing laggard chromosomes (Fig. 4B,D). Thus, lagging chromosomes were probably the origin of the micronuclei observed at the later tetrad stage. No splitting of sister chromatids or chromosome fragmentation such as was previously reported in the CRISPR *Tazip4-B2* mutant [26], was observed in the *zip4-ph1d* mutant. These observations were consistent with our previous observation that pairs of univalents were frequently close to each other and oriented to different poles of the nucleus. This suggests that they could have been connected to each other as they orientated on the metaphase I plate, which might then have facilitated correct segregation of these univalents.

### The *zip4-ph1d* mutation allows homoeologous crossover formation in wheat haploids

Previous studies revealed that in polyploid wheat, *TaZIP4-B2* is involved in both promotion of homologous pairing-synapsis, and suppression of homoeologous CO [17,25]. Accordingly, a CRISPR *Tazip4-B2* mutant exhibited disrupted pairing-synapsis, whilst allowing CO between homoeologues. However, the present study indicates that in the *zip4-ph1d* mutant, carrying the TaZIP4-B2 with a A^167^V substitution, promotion of homologous pairing-synapsis is preserved, as chiasmata still occur between pairs of homologues. This then raised the question as to whether the TaZIP4-B2 in the *zip4-ph1d* mutant still possessed the ability to suppress homoeologous CO. In a previous study, hybrids between the *zip4-ph1d* mutant and *Ae. variabilis* showed an increase in homoeologous CO, similar to that found when the original Sears 59.3Mb *ph1b* deletion mutant was crossed with this wild relative [20,25]. This suggests that the *zip4-ph1d* mutant carries a partial ‘loss-of-function’ *TaZIP4-B2*, which has lost the capability to suppress homoeologous CO between wheat and its wild relatives, whilst retaining the ability to promote homologous pairing and synapsis. However, it had been suggested that *Ae. variabilis* could also be contributing to the induction of homoeologous CO observed in these hybrids. Therefore, it was important to determine whether the partial ‘loss-of-function’ *TaZIP4-B2* copy carried by the *zip4-ph1d* mutant still suppressed homoeologous CO between wheat chromosomes when there were no wild relative chromosomes present.

To clarify this situation, we obtained several independent haploids of both the *zip4-ph1d* mutant and wild type Cadenza wheat. GISH was performed on metaphase root tip spreads from both sets of haploids, to confirm the correct number and composition of chromosomes. A complete haploid set of 21 homoeologous wheat chromosomes was present in all haploids, each composed of three sets of 7 chromosomes corresponding to the three ancestral genomes A, B and D (Fig. 5A). In such haploids, each A genome homoeologue for instance, has the potential to synapse and CO with a B or D genome homoeologue, and vice versa. We scored meiotic configurations at metaphase I in three haploids from each of the *zip4-ph1d* mutant and wild type wheat. This analysis revealed an average of more than six chiasmata (6.26) per meiocyte in the *zip4-ph1d* mutant haploids and less than one chiasmata (0.63) per meiocyte in the Cadenza wild type haploids (Fig. 5B,C; Table 2; Supplementary Table S2). Levels of rod bivalents and chiasmata observed in the *zip4-ph1d* mutant haploids were similar to meiotic scores previously reported for haploids generated from the Sears 59.3Mb *ph1b* deletion mutant [31]. This data confirms that the effect of the A^167^V substitution in *TaZIP4-B2* carried by the *zip4-ph1d* mutant is sufficient to allow CO between homoeologues in wheat haploids, to the same level as observed in haploids derived from the *ph1b* mutant. The reduced level of chiasmata observed in the wild type Cadenza haploid, compared to that seen in the *zip4-ph1d* mutant and Sears 59.3Mb *ph1b* mutant haploids, conclusively confirms that the wild type *TaZIP4-B2* gene on chromosome 5B suppresses COs between homoeologous wheat chromosomes.

**Figure 5.**
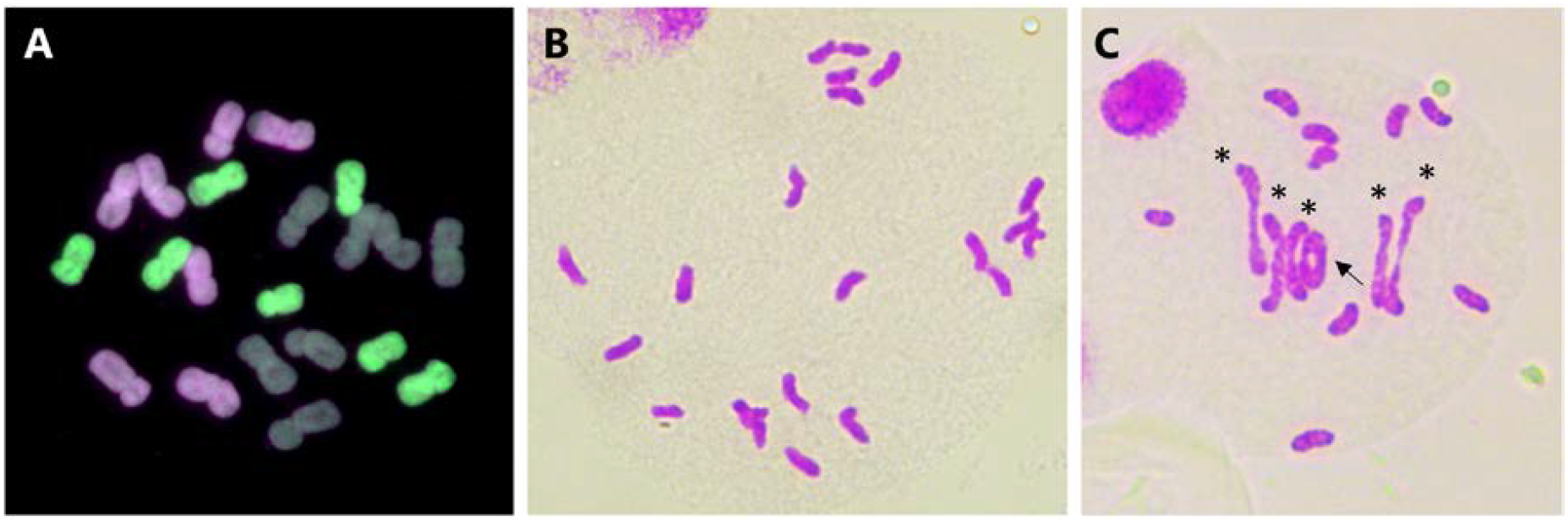
Partial loss-of-function *TaZIP4-B2* mutation (*zip4-ph1d*) allows CO formation between homoeologous chromosomes in haploid wheat. (**A**) Root-tip metaphase of one representative of the haploids (n = 3x = 21) labelled by genomic *in situ* hybridisation (GISH). Seven A-genome chromosomes are shown in magenta, seven B-genome chromosomes in grey and seven D-genome chromosomes are shown in green. (**B, C**) Representative meiotic metaphase I configurations of haploid Cadenza WT (**B**) and haploid *zip4-ph1d* (**C**) stained by Feulgen. Haploid Cadenza WT shows no chiasma while haploid *zip4-ph1d* displays five rod bivalents (indicated by an asterisk) and one ring bivalent (indicated by an arrow).

**Table 2.** Frequencies of rod bivalents, ring bivalents, multivalents and total chiasmata in the haploid Cadenza wild type (Cad-WT) and haploid Cadenza *zip4-ph1d* mutant. Mean number per cell and range of variation between cells are indicated.

| Line | No. of cells examined | Rod bivalents |  | Ring bivalents |  | Multivalents |  | Chiasma frequency per cell |  |
| --- | --- | --- | --- | --- | --- | --- | --- | --- | --- |
| | | Mean ± SE | Range | Mean ± SE | Range | Mean ± SE | Range | Mean ± SE | Range |
| Haploid Cad-WT | 566 | 0.60 ± 0.04 | 0-3 | 0.01 ± 0.003 | 0-1 | 0.005 ± 0.005 | 0-1 | 0.63 ± 0.03 | 0-5 |
| Haploid *zip4-ph1d* | 433 | 3.36 ± 0.04 | 0-6 | 0.66 ± 0.07 | 0-4 | 0.68 ± 0.08 | 0-3 | 6.26 ± 0.12 | 0-12 |

### Homoeologous crossover suppression by *ZIP4* is not essential to maintain wheat fertility

In the CRISPR *Tazip4-B2* mutant, both the ‘promotion of homologous pairing-synapsis’ and the ‘homoeologous CO suppression’ phenotypes are lost [26]. As a result, 56% meiocytes exhibit major meiotic abnormalities: 26% meiocytes have univalents, and 32% show both univalents and multivalents. This leads to 50% pollen grains being smaller, and a subsequent 50% reduction in grain number. Conversely, most meiocytes in the *zip4-ph1d* mutant exhibit regular CO formation between homologues leading to correct chromosome segregation. As a result, only 12.2% tetrads exhibit micronuclei, and there is less than 15% small pollen. The grain number in the *zip4-ph1d* mutant is therefore the same as that found in wild type Cadenza, despite CO still occurring between wheat homoeologues in haploids derived from the mutant.

Thus, the present study shows that homoeologous CO suppression by *TaZIP4B2* may not be essential for wheat fertility if promotion of homologous pairing-synapsis is still functioning. These observations are similar to those recently reported, where homoeologous CO was induced in wheat-wild relative hybrids through the mutation of the *Ph2*/*TaMSH7-3D* gene, but this did not result in a significant reduction in fertility [34]. In both cases, the promotion of homologous pairing-synapsis by *TaZIP4-B2* would still be functioning. These observations suggest that promotion of homologous pairing-synapsis may be more important for prevention of major meiotic abnormalities and reduced fertility in polyploid wheat, than it is suppression of homoeologous CO. This promotion of homologous pairing-synapsis appears to account for the majority of the 50% preservation of grain number by *TaZIP4-B2*. Thus, on the polyploidisation of wheat, and despite the pre-existence of three *ZIP4* copies, duplication and divergence of *TaZIP4-B2* was required to promote homologous pairing-synapsis and prevent a potential 50% loss of grain number. These observations contrast with previous studies of polyploid brassicas, which exhibited more extensive chromosome rearrangements and loss of gene content upon polyploidisation [35,36] than is observed in wheat [9]. The absence of the main locus limiting homoeologous recombination in brassicas, *BnaPh1* [37], reveals the importance of homoeologous CO suppression for preservation of genome stability in other polyploids. This suggest that different polyploid species have reached genome stability through different evolutionary mechanisms.

*ZIP4* is required for pairing, synapsis and CO in budding yeast [23,24]. However, in Arabidopsis and rice, *ZIP4* is only required for CO, and not for pairing and synapsis [21,22]. Therefore, it is unlikely that the promotion of homologous pairing-synapsis by *TaZIP4-B2* in 56% of wheat meiocytes is a direct effect. When *TaZIP4-B2* is absent, homologous chromosome sites are more dispersed in the nucleus during the telomere bouquet [15,38,39]. Thus, it is possible that in wheat, the earlier and higher expression of *TaZIP4-B2*, compared to the later and lower expression of the group 3 *TaZIP4* copies [13], enables the TaZIP4-B2 protein to be loaded earlier onto chromosomes, altering their conformation, and reducing the dispersion of the homologous sites. This in turn, might enable more rapid pairing and synapsis of the homologous sites during the telomere bouquet. Altered chromosome conformation in the presence of the TaZIP4-B2 protein would be consistent with the observations in rice, where the loss of *ZIP4* results in a slight decondensation of meiotic chromosomes [22]. Duplication and divergence of *TaZIP4-B2* must have led to the promotion of the pairing-synapsis function being rapidly acquired, given its importance in preventing major abnormality and reduced fertility, and the lack of extensive chromosome rearrangement or gene loss in wheat.

Variable levels of homoeologous CO have been reported in crosses of different polyploid Brassica genotypes [35,36, 37]. Interestingly, it has also been reported that wild tetraploid wheat (*Triticum turgidum* ssp dicoccoides) exhibits variable levels of homoeologous CO in crosses with a wild relative. This variation corresponds with different geographical locations in Israel and therefore may have had adaptive value [40]. This may suggest that the ‘homoeologous CO suppression’ phenotype evolved through divergence of *TaZIP4-B2* over a long time period and may have been associated with adaptation to different environments. It would be interesting to explore whether there is an association between homoeologous CO frequencies involving different allelic variations of *TaZIP4-B2* in different wheat varieties, and the varying environmental conditions found in different geographical regions.

## Conclusions

*TaZIP4-B2* plays two complementary roles to ensure wheat genome stability and fertility upon polyploidisation: promotion of homologous pairing-synapsis and suppression of homoeologous CO. This results in major meiotic abnormalities and 50% loss of grain number in the CRISPR *Tazip4-B2* and *ph1b* mutants, where both functions are lost. The present study analyses the *zip4-ph1d* mutant carrying a *Tazip4-B2* mutation where only the ‘suppression of homoeologous CO’ function is lost, but the ‘promotion of homologous pairing-synapsis’ is still functional, allowing us to separate both functions for the first time. Remarkably, this *zip4-ph1d* mutant shows mostly balanced chromosome segregation and wild type fertility, confirming that loss of homoeologous CO suppression between wheat chromosomes does not in itself result in major meiotic abnormality or reduced fertility, when promotion of homologous pairing-synapsis by *TaZIP4-B2* is still functional. Thus, when impact on fertility for agriculture and human nutrition are considered, the effect of *TaZIP4-B2* on promotion of homologous pairing and synapsis appears more important than its suppressant effect on homoeologous CO. However, the ‘loss of homoeologous CO suppression’ phenotype is equally important in wheat breeding, as manipulation of this phenotype through use of the *ph1b* mutant has been exploited extensively to introgress advantageous chromosome segments from wild relatives, saving the global economy billions of dollars over the years in this major crop. We therefore suggest that the *zip4-ph1d* mutation in an elite wheat background should now be used in breeding for this purpose, as in contrast to the *ph1b* mutant, it will not accumulate such extensive rearrangements and will not exhibit reduced fertility.

## Materials and Methods

### Plant material

The *zip4-ph1d* mutant used in this work was formerly designated as *Tazip4-B2-*Cadenza1691 in Rey et al, 2017 [20]. This mutant is a hexaploid wheat *T. aestivum* cv. Cadenza TILLING mutant (Cadenza1691) carrying a single missense mutation (C to T change at the position 500 in the CDs, leading to an amino acid substitution A^167^V in the SPO22 domain) [20]. The mutant was backcrossed with Cadenza three times to clean the background of the TILLING mutations, and the SPO22 missense mutation selected each time using KASP markers (Hex: GGGCGATCAGTCCCTCGC; Fam: GGGCGATCAGTCCCTCGT; Common: ATCTGGTATACTTGCGGGGC; KASP 2X Master Mix, LCG, Middlesex, UK), following the manufacturer’s instructions.

All plant material was grown in a controlled environmental room under the following growth conditions: 16 h/8 h, light/dark photoperiod at 20°C day and 15 °C night, with 70% humidity.

### Pollen profiling

Ten plants for the *zip4-ph1d* mutant and wild type were included in this experiment. Mature yellow anthers (just before shedding pollen) were collected in 0.5 mL 70% ethanol, from five main florets at the middle portion of the first spike of each plant as described previously [26]. Pollen grains were released from anthers by sonication. Size and number of filtered pollen grains were measured using a Coulter counter (Multisizer 4e, Beckman Coulter Inc.), fitted with a 200 µm aperture tube [26]. Pollen number distribution and analysis were performed using an R script to calculate plot differential pollen size distribution and pollen number per anther as described previously [26].

### Pollen viability

Pollen viability was assessed using Alexander stain [41]. Fresh wheat pollen grains from three anthers were shed on a droplet of Alexander stain placed on a microscope slide and images covering the whole slide were taken for scoring. Three biological replicates, each with >1000 pollen grains were analysed for each genotype.

### Grain setting assessment

The first three bagged spikes from each plant were harvested when fully dried and threshed separately after counting spikelet number. Number of grains per spike was then measured using the MARVIN grain analyser (GTA Sensorik GmbH, Neubrandenburg, Germany). Grain setting was then calculated as a normalized grain number per spike ((actual grain number per spike/expected grain number per spike)*100), in order to eliminate the effect of different number of spikelets per spike on grain number. Expected grain number per spike was calculated by multiplying number of spikelets by three, considering that each spikelet has three main fertile florets.

### Haploid production

Haploid production of the *zip4-ph1d* mutant and Cadenza wild type was performed as previously described [42] with some modifications. Briefly, all plant material was grown in a controlled environmental room as described in the plant material section. A solution of 2,4-D 5mg/l + AgNO3 100 mg/l was applied in vivo by filling each floret with the solution, 24h after pollination with maize. Wheat regeneration media described in Hayta et al, 2021 [43], was used to culture the embryo rescued. Ploidy level of the seedlings was determined using GISH (genomic *in situ* hybridisation) in metaphase I spreads from root tips as previously described [44,45]. *Triticum urartu*, and *Aegilops tauschii* were used as probes to label wheat A- and wheat D-genomes, respectively. *Aegilops speltoides* genomic DNA was used as a competitor in the hybridisation mix. *T. urartu*, and *Ae. tauschii* genomes were labelled with biotin-16-dUTP and digoxigenin-11-dUTP using the Biotin- or the DIG-nick translation mix, respectively, according to the manufacturer’s instructions (Sigma, St. Louis, MO, USA). Biotin-labelled probes were detected with Streptavidin-Alexa 660 (Thermo Fisher Scientific, Waltham, MA, United States); digoxigenin-labelled probes were detected with anti-digoxigenin-FITC (Sigma, St. Louis, MO, USA).

### Meiotic analysis

Anthers at the right meiotic stage were collected as previously described [15], fixed in 100% ethanol/acetic acid 3:1 (v/v) and kept at 4 °C until needed. Cytological analysis of Pollen Mother Cells using the Feulgen technique has been described elsewhere [46]. GISH on meiotic metaphase I spreads was performed as described in Rey et al, 2021 [45]. *T. urartu*, and *Ae. tauschii* were used as probes to label wheat A- and wheat D-genomes respectively, as described above in the haploid production section. *Ae. speltoides* genomic DNA was used as a competitor in the hybridization mix.

### Image processing

Pollen Mother Cells stained by the Feulgen technique were imaged using a LEICA DM2000 microscope (Leica Microsystems, http://www.leica-microsystems.com/), equipped with a Leica DFC450 camera and controlled by LAS v4.4 system software (Leica Biosystems, Wetzlar, Germany).

All cells labelled by GISH were imaged using a Leica DM5500B microscope equipped with a Hamamatsu ORCA-FLASH4.0 camera and controlled by Leica LAS X software v2.0. Images were processed using Adobe Photoshop CS5 (Adobe Systems Incorporated, US) extended version 12.0 × 64.

### ZIP4 protein sequence analyses

Protein sequences of the TaZIP4 genes and their orthologs were retrieved from the Ensemble database (Table S4). Multiple protein sequence alignment was performed using the Clustal X programme (version 2) [47,48]. TPR domains were predicted using the TPRpred program (https://toolkit.tuebingen.mpg.de/tools/tprpred) [49]. A graphical representation of the conserved residues in the aligned protein sequences was produced using the Skyling webtool (https://skylign.org/logo/) [50]. The secondary structure (2D) of the TPR2 domain in the ZIP4 proteins was predicted using the PSSM random forest based model in the PEP2D server (Prediction module of PEP2D (osdd.net)) [51], and using the Jpred 4 protein secondary structure prediction server (www.compbio.dundee.ac.uk/jpred) [52].

## Author Contributions

ACM produced the haploids for the *zip4-ph1d* and Cadenza wild type mutants. ACM carried out the cytological experiments and analysis of the produced data. AKA undertook the pollen and grain set analysis on the *Tazip4-B2* mutant and the ZIP4 protein sequence analyses. GM provided thoughts and guidance and revised and edited the manuscript produced by ACM and AKA. All authors have read and agreed to the published version of the manuscript.

## Additional information

### Data Availability Statement

The data that supports the findings of this study are available in the supplementary material of this article and from the corresponding author upon reasonable request.

