## Supplemental Material for "A separation-of-function *ZIP4* wheat mutant allows crossover between related chromosomes and is meiotically stable"

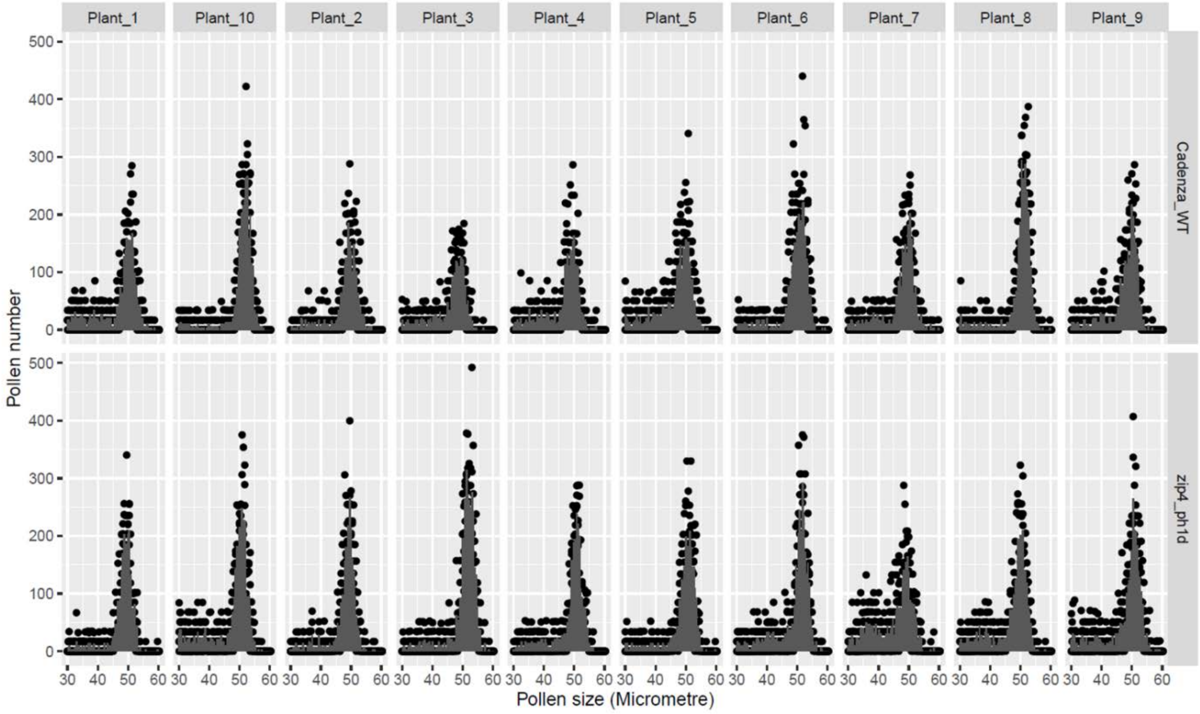


**Supplementary Figure S1.** The differential pollen size distribution data collected by coulter counter (Multiziser 4e) for each plant (biological replicate) from the *zip4-ph1d* mutant and its corresponding wild type (cv. Cadenza). Five anther samples were collected from florets in the middle portion of the first spike of each plant. Each sample contains three anthers collected from the same floret. Pollen profiles are consistent within the same genotype and very similar in both genotypes.

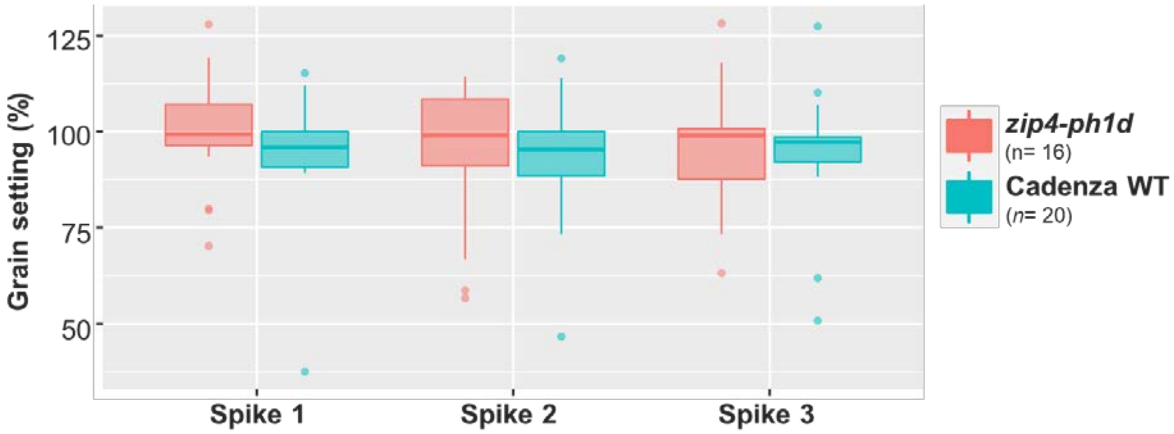


**Supplementary Figure S2.** Grain setting (normalized grain number per spike) in the first three spikes of the plant for the *zip4-ph1d* mutant and the wild type control (cv. Cadenza) under CER growth conditions. *n* refers to the number of plants (biological replicates) used in the experiment.

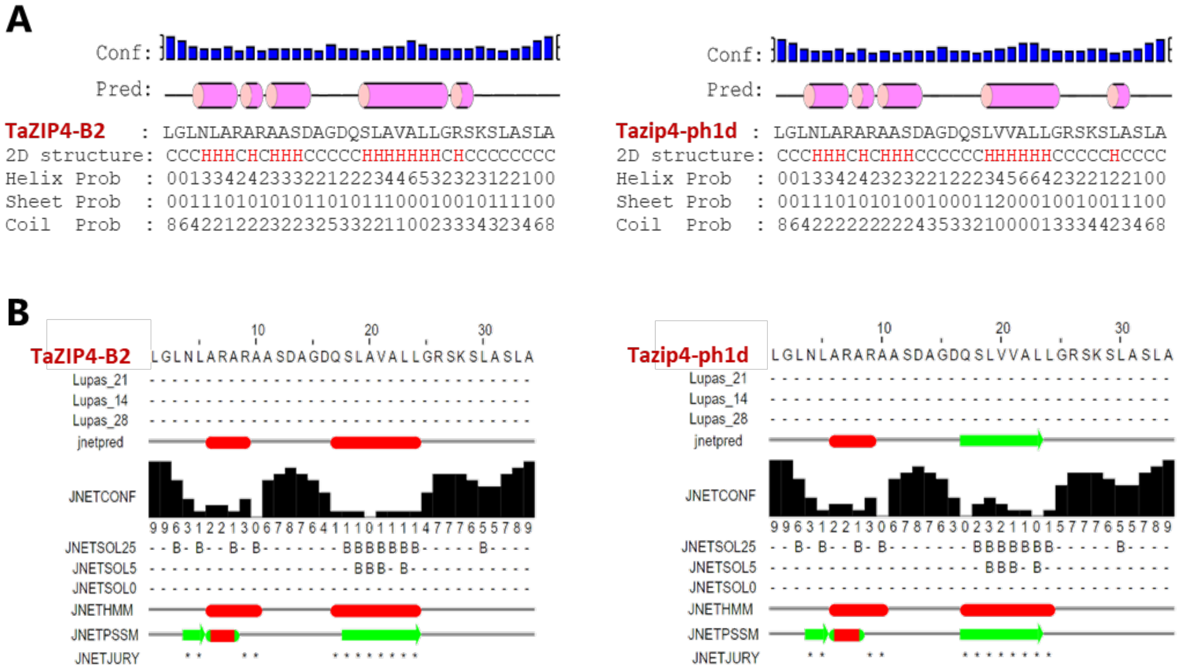


**Supplementary Figure S3. Prediction of the secondary structure of the TPR2 of in the wild type TaZIP4-B2 protein and mutant (Tazip4-ph1d).** (**A**) 2D structure prediction using the PEP2D server based on PSSM random forest model. Helix is represented in cylindrical shape in pink colour and coil is represented as black line. (**B**) 2D structure prediction using the Jpred 4 server. Key: Jnet: Final secondary structure prediction for query; Jalign: Jnet alignment prediction; Jhmm: Jnet hmm profile prediction; Jpssm: Jnet PSIBLAST pssm profile prediction; Lupas: Lupas Coil prediction (window size of 14, 21 and 28). Jnet_25: Jnet prediction of burial, less than 25% solvent accessibility; Jnet_5: Jnet prediction of burial, less than 5% exposure; Jnet_0: Jnet prediction of burial, 0% exposure; Jnet Rel: Jnet reliability of prediction accuracy, ranges from 0 to 9, higher scores indicate greater significance.

**Supplementary Table S1.** Analysis of the level of micronuclei (MN) at the tetrad stage in (**A**) Cadenza wild type (WT) and (**B**) the *zip4-ph1d* mutant. Three individual plants were used from each genotype.

**A**

| | Cadenza WT-1 | Cadenza WT-2 | Cadenza WT-3 | All Cadenza WT |
|---|---|---|---|---|
| **% Tetrads with MN** | 8.02 | 4.50 | 2.91 | 5.49 |
| **Total No. Tetrads** | 324 | 111 | 275 | 710 |
| **No. Tetrads with MN** | 26 | 5 | 8 | 39 |

**B**

| | *zip4-ph1d*-1 | *zip4-ph1d*-2 | *zip4-ph1d*-3 | All *zip4-ph1d* |
|---|---|---|---|---|
| **% Tetrads with MN** | 10.52 | 13.91 | 11.82 | 12.26 |
| **Total No. Tetrads** | 190 | 273 | 279 | 742 |
| **No. Tetrads with MN** | 20 | 38 | 33 | 91 |

**Supplementary Table S2.** Frequencies of rod bivalents, ring bivalents, multivalents and total chiasmata in three haploid Cadenza wild type (Cad-WT) and three haploid Cadenza *zip4-ph1d* mutant plants. Mean number per cell and range of variation between cells are indicated.

**A**

| Line | No. of cells examined | Rod bivalents | | Ring bivalents | | Multivalents | | Chiasma frequency per cell | |
|---|---|---|---|---|---|---|---|---|---|
| | | Mean | Range | Mean | Range | Mean | Range | Mean | Range |
| **Haploid Cad-WT-1** | **135** | 0.585 | 0-3 | 0.007 | 0-1 | 0.015 | 0-1 | 0.630 | 0-5 |
| **Haploid Cad-WT-2** | **158** | 0.677 | 0-3 | 0.006 | 0-1 | 0.000 | 0 | 0.690 | 0-3 |
| **Haploid Cad-WT-3** | **273** | 0.549 | 0-2 | 0.015 | 0-1 | 0.000 | 0 | 0.579 | 0-3 |

**B**

| Line | No. of cells examined | Rod bivalents | | Ring bivalents | | Multivalents | | Chiasma frequency per cell | |
|---|---|---|---|---|---|---|---|---|---|
| | | Mean | Range | Mean | Range | Mean | Range | Mean | Range |
| **Haploid *zip4-ph1d-1*** | 104 | 3.288 | 0-6 | 0.760 | 0-4 | 0.788 | 0-3 | 6,423 | 2-10 |
| **Haploid *zip4-ph1d-2*** | 102 | 3.363 | 0-6 | 0.529 | 0-4 | 0.529 | 0-3 | 6.020 | 1-11 |
| **Haploid *zip4-ph1d-3*** | 227 | 3.441 | 0-6 | 0.696 | 0-3 | 0.718 | 0-3 | 6.326 | 0-12 |
